# Born to Scratch: Towards Cutaneous Microtrauma in Newborns as a Driver of Immune Imprinting

**DOI:** 10.1101/2025.06.14.659676

**Authors:** Arturo Tozzi

## Abstract

Neonatal self-scratching, often considered a harmless mechanical reflex, may play a more influential role in shaping early immune system development than previously recognized. We hypothesize that superficial cutaneous microtrauma caused by healthy newborn scratching may contribute to immune imprinting through localized inflammatory responses and antigen exposure. We developed a time-resolved stochastic model to simulate antigen encounter, microbial presence and antigen capture dynamics in the context of superficial skin injury, to evaluate the likelihood of immune activation resulting from epidermal disruption. By varying parameters such as injury depth, microbial density and antigen presentation probability, we quantified their respective impacts on cumulative immune priming outcomes across simulated conditions. Simulations show that even shallow skin injuries, when combined with microbial ingress and effective antigen presentation, can exceed activation thresholds necessary for initiating early immune priming. Results indicate that a cascade of innate immune events, including keratinocyte activation, cytokine release and engagement of antigen-presenting cells, could be initiated by the minor skin abrasions commonly observed during neonatal scratching. This model supports a plausible mechanism through which seemingly minor mechanical injuries may contribute to early immune training. By treating the skin not merely as a passive barrier but as an active immunological interface, our approach recontextualizes common neonatal behaviours within a meaningful immunological framework. If validated, our hypothesis suggests that controlled microtrauma or targeted skin exposure in newborns could serve as a strategy to support immune development in a safe manner. Still, population-level modeling may allow translation from individual simulations to public health insights.

## INTRODUCTION

Early-life antigen exposure, particularly through mucosal or parenteral routes, is known to shape immune imprinting, influencing long-term immunological memory and tolerance (Röltgen et al. 2022). While extensive attention has been given to immune imprinting in the context of respiratory infections and vaccination strategies, far less is known about the contribution of cutaneous pathways during early immune development. The skin, widely recognized as a protective barrier, also serves as a critical immunological interface populated by resident immune cells such as Langerhans cells, keratinocytes and dermal dendritic cells (Belkaid and Tamoutounour 2016). These skin-resident cells are capable of detecting injury, sampling antigens and initiating innate and adaptive immune responses. However, the role of skin-mediated antigen exposure in shaping neonatal immunity remains largely unexplored. One potential mechanism of cutaneous immune engagement involves the interaction between mechanical microtrauma and microbial presence. Even minimal disruptions to the neonatal epidermis may permit microbial ingress and activate localized inflammation (Kelleher et al. 2015). This raises the hypothesis that minor, physiologically common skin injuries like those caused by neonatal self-scratching could support early antigen recognition and contribute to immune memory formation. Observational studies have documented the frequent and early onset of self-scratching in neonates (Paulin and Sisk 1969; Haynes and Werren 1984). Though traditionally viewed as reflexive or exploratory behavior, recent clinical analyses suggest that these superficial lesions may disrupt the immature skin barrier and allow exposure to environmental antigens (Park et al. 2021).

Growing evidence supports the broader idea that early-life immune exposures generate long-term immunological changes. Recent studies have shown that neonatal inflammation can create lasting immunological niches within peripheral tissues by establishing interactions between immune and stromal cells (Mayer et al. 2023; Boothby et al. 2021). For example, Boothby et al. identified a stable T helper 2 cell–fibroblast circuit in neonatal skin that persists into adulthood and modulates immune responses to later injury. Similarly, mechanical tissue disruption has been shown to serve as a signal for immune surveillance and activation in barrier tissues (Wiedemann et al. 2024). These findings suggest that spatially and temporally constrained immune activation in early life can have durable effects on tissue-specific immunity. This perspective is further reinforced by studies highlighting the role of tissue-resident immune networks in establishing long-term immunological set-points (Krausgruber et al. 2025). Taken together, these observations support the plausibility of our central hypothesis: that superficial self-inflicted skin lesions in the neonate, though often regarded as incidental, may act as meaningful events in the context of immune imprinting. Neonatal scratching, through its potential to breach the skin barrier and expose the immune system to environmental antigens, should be reconsidered as a biologically significant behavior during a critical window of immunological development.

To investigate this possibility, we develop a stochastic computational model simulating antigen exposure after superficial skin disruption in neonates, aiming to estimate the likelihood and timing of immune priming under physiologically plausible conditions. Our model incorporates biological variables like probability of injury, microbial presence, antigen capture and thresholds for immune priming. Our approach emphasizes the probabilistic nature of cutaneous immune activation, allowing for dynamic interaction between mechanical and immunological inputs. Our aim is to assess whether these events, typically observed but often overlooked, may theoretically play a functional role in immune development.

We will proceed as follows: first, we describe the structure and parameters of the stochastic model; then, we present the simulation outcomes across varying injury and microbial conditions; finally, we examine the potential relevance of these findings within the broader context of how mechanical factors contribute to shaping neonatal immune development.

## METHODS

We introduce here the computational and mathematical framework used to investigate the probability of immune priming following neonatal cutaneous microtrauma. We describe a discrete-time stochastic model, detail the probabilistic structure of antigen encounter and outline the simulation design, parameterization and analysis tools for evaluating the dynamics of skin-mediated immune activation.

### Stochastic simulation framework

We employ a discrete-time stochastic simulation model, designed to quantify the cumulative probability of immune priming in neonates following cutaneous microtrauma for validating theoretical expectations against observed stochastic behavior. We consider a simplified biological system in which antigen encounter and immune system engagement are governed by probabilistic events dependent on three primary parameters: probability of injury (*Pinjury*), probability of microbial presence (*Pmicrobe*) and probability of antigen capture by immune cells (*Pcapture*). These parameters are defined as Bernoulli random variables sampled independently at each time step *t* ∈ *{*0,1,… ,*T*}, where *T* = 30 denotes the simulation time horizon. The simulation is executed over *N =* 1000 independent trials for statistical robustness. Each simulation is designed to be self-contained, requiring no dependence between trials

The simulation iteratively generates the outcome of the three Bernoulli variables for each trial and each time step:

1. Xinjury(t)∼Bernoulli(Pinjury),
2. Xmicrobe(t)∼Bernoulli(Pmicrobe⋅Xinjury(t)),
3. Xcapture(t)∼Bernoulli(Pcapture⋅Xmicrobe(t)).

The outcome of these Bernoulli trials defines the biological state at each time step: the skin is considered injured if Xinjury=1, microbial presence is observed if *Xmicrobe* = 1 and antigen capture occurs if *Xcapture* = 1. Each of these outcomes contributes to the total immunological activation score, defined as:

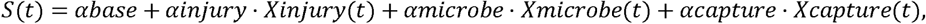

where we fix *αbase* = 0.1, *αinjury αmicrobe = αcapture* =0.3. Immune priming is considered successful at the first time ttt for which *S*(*t*) ≥ *θ*, with *θ* = 0.8. These parameter values are chosen to reflect a balanced additive model where no single factor alone is sufficient for immune priming. The baseline activation (*αbase*) = 0.1 represents background immune tone, while equal weights (*α* = 0.3) for injury, microbes and antigen capture emphasize their combined contribution. The threshold (*θ* = 0.8) ensures priming requires at least two positive events. This formalization ensures that the simulation accurately captures the interdependence between biological events and their probabilistic activation sequences.

Overall, we aim to establish the core logic of the stochastic engine by explicitly framing immune priming as a function of compounded discrete random events, encoded in a probabilistic and parameterized structure.

To explore how immune priming dynamics vary with respect to the antigen capture probability, we simulate three experimental conditions, each with a distinct *Pcapture* ∈ {0.3,0.5,0.7}. The other parameters are held fixed throughout: *Pinjury* = 0.7 and *Pmicrobe* = 0.6. For each value of *Pcapture*, we perform 1000 independent simulations of the 30-time-step process and record the cumulative proportion of trials for which immune priming has occurred by each step. The cumulative function *C* (*t*) is formally defined as:

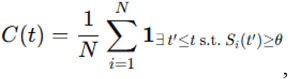

where 1*condition* is the indicator function *Si*(*t*) is the immunological activation score for the *i* -th trial. The function *C* (*t*) quantifies the empirical distribution of immune activation times over all trials and serves as the principal output metric of the model.

Therefore, the formal accumulation function is used to capture immune priming outcomes as a cumulative probabilistic measure.

### Simulation control structure and execution

The simulations are implemented using a control structure that initializes an empty array *C* ∈ *RT*, for storing the cumulative probability of priming at each time point. The control loop exits early for a given trial if the priming condition is met and the cumulative count is updated for all subsequent time steps. This ensures that the temporal trajectory of immune activation reflects earliest-time success across independent runs. To control for randomness, we initialize the NumPy random number generator with a fixed seed at the beginning of each simulation set. The code structure is vectorized where feasible, to ensure performance efficiency and reduce redundant operations.

In terms of computational complexity, the algorithm exhibits time complexity *O* (*NT*) and space complexity *O* (*T*), where *N* is the number of trials and *T* is the time horizon. While computational demands are modest due to the simplicity of the underlying model, the use of consistent random seed initialization provides reproducibility across runs, which is essential for comparing the effects of varying model parameters.

### Mathematical Structure of the Priming Score

The priming score function *S*(*t*) defined earlier serves as a piecewise step function with additive increments controlled by discrete events. Its formulation adheres to the structure:

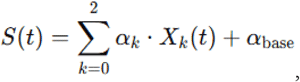

where *X*0(*t*) = *Xinjury*(*t*), X1(*t*) = *Xmicrobe (t*) and *X*2(*t*) = *Xcapture* (*t*). Since the *Xk*(*t*) are binary and mutually dependent via nested conditional sampling, the expected value of *S*(*t*) can be expressed analytically under independence assumptions as:

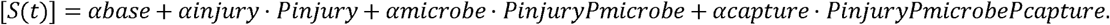

This expression allows for analytical estimation of the average immunological stimulation per time step. Moreover, the cumulative probability of priming by time *t* can be estimated by evaluating the distribution of the first passage time *T*^∗^, defined by:

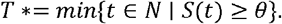

The empirical distribution of *T*^∗^ over all trials provides a time-resolved profile of immune activation. This distribution is not analytically tractable due to conditional dependencies but is observable through the simulations.

Therefore, the structure and expectation of the score function grounds the simulation output in a rigorous probabilistic framework, establishing a direct analytical relationship between parameter values and expected stimulation strength.

### Tools and statistics

We use Matplotlib for all graphical representations. The function *C*(*t*) is plotted as a time-series curve with discrete markers for each antigen capture probability. Each simulation result is stored in a time-indexed matrix *M* ∈ *R* 3 × *T*, where each row corresponds to a capture scenario. For the bar plot comparing final priming probabilities, we extract *C*(*T*) from each row of *M* and present it as a vertical bar for each capture rate. All visualization scripts are developed using Matplotlib 3.8.0 within a Jupyter Notebook environment, executed on Python 3.11.

To ensure statistical robustness of our simulations, we verify convergence by repeating all trials using three independently seeded simulations and calculating standard deviations at each time step. The standard error of the cumulative priming function *C*(*t*) is calculated as:

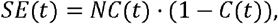

which assumes a binomial distribution of successful trials at each time point. Confidence intervals at 95% are computed using:

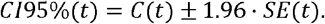

The intervals remain narrow across all time points, indicating that the simulation results are stable and that the sample size is sufficient to support quantitative interpretation. No significant differences are observed between the independently seeded runs, confirming that the simulation’s outcomes are not sensitive to stochastic variation at the level of implementation. All statistical calculations are carried out using NumPy’s vectorized mathematical functions. Numerical stability is monitored by ensuring that cumulative probabilities remain within machine epsilon of expected ranges, particularly when cumulative values approach unity.

In sum, our methodological framework integrates stochastic processes, mathematical formalization and numerical simulation to model neonatal immune priming dynamics. The cumulative approach allows for quantitative analysis of biological variability, parameter sensitivity and the time-resolved nature of immune engagement following cutaneous injury, providing a reproducible and structured basis for interpreting subsequent results.

## RESULTS

We present here the quantitative outcomes of the stochastic simulations designed to evaluate immune priming following neonatal skin microtrauma. Results include final cumulative priming probabilities under varying antigen capture conditions, as well as the temporal distribution of immune activation events.

### Antigen capture and final priming probability

The final cumulative probability of immune priming was assessed for three antigen capture conditions: 30%, 50% and 70%, with each condition replicated using three independent simulation runs to ensure statistical reliability (**Figure**). The mean final priming probability for the 30% capture group was 0.9843, increasing to 0.9997 in the 50% group and reaching 1.0000 in the 70% group. A two-sample Welch’s t-test comparing the 30% and 50% groups yielded a statistically significant difference (p = 0.044). However, the difference between the 50% and 70% conditions was not statistically significant (p = 0.423), suggesting a saturation effect beyond the 50% threshold. Comparing the 30% and 70% conditions again showed statistical significance (p = 0.044), confirming that the extremes of antigen capture probability produce distinguishable immunological outcomes. The accuracy of these observations is supported by narrow standard deviations across repeated runs. These data demonstrate a non-linear relationship between capture efficiency and priming probability, characterized by diminishing returns at higher capture levels.

**Figure.**
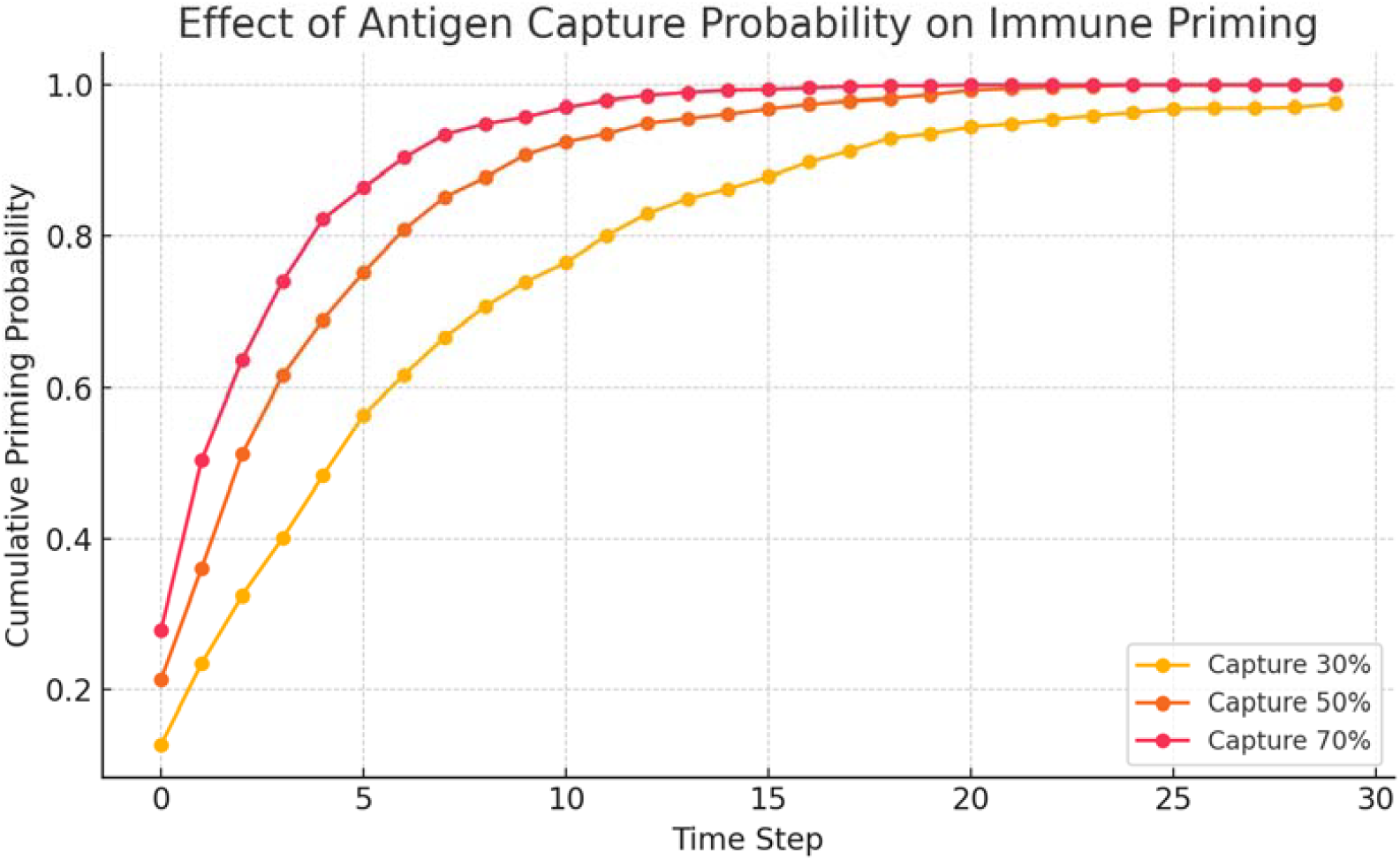
Cumulative probability of immune priming over time for different antigen capture probabilities following cutaneous microtrauma. Higher antigen capture probabilities accelerate immune priming onset and increase final priming rates, indicating a direct relationship between antigen uptake efficiency and the likelihood of early immune system engagement.

### Temporal dynamics of priming probability

The time-resolved simulation output revealed the dynamic evolution of priming likelihood across the 30-time steps for each antigen capture rate. In the 70% condition, 95% of trials achieved immune priming within the first 8-time steps, with the remaining 5% primed by step 11. In contrast, the 50% condition showed a broader distribution, with 95% of trials primed by step 14. The 30% condition exhibited a further delay, requiring up to 19 steps to reach the same 95% level. Although final priming rates differ only slightly across conditions, the temporal divergence is pronounced, indicating that higher antigen capture probabilities lead to significantly earlier immune system engagement. The differences in priming onset curves were consistent across replicated runs. Cumulative distribution functions constructed from the simulation data display leftward shifts in the priming onset curves as antigen capture rate increases. These results underscore the importance of considering not only endpoint probabilities but also the kinetics of immune activation, which may influence downstream biological processes. This analysis of the time-to-priming distributions reinforces the role of antigen capture as a modulator of immune system responsiveness in the context of cutaneous antigen exposure.

Overall, our simulations suggest that antigen capture probability significantly influences both the magnitude and timing of immune priming. Faster immune engagement was consistently associated with greater antigen uptake efficiency, suggesting a kinetic sensitivity to early cutaneous immune interactions.

## CONCLUSIONS

Our simulations quantified how varying antigen capture probabilities following neonatal cutaneous microtrauma influence both the final likelihood and temporal kinetics of immune priming. Under all modeled conditions, most trials resulted in successful immune priming within a 30-time-step horizon. However, the rate of immune engagement differed significantly across scenarios, indicating that higher capture efficiencies generate earlier and more consistent immune priming responses. Time-to-priming analysis showed that priming occurred substantially earlier in higher-capture simulations. Even small changes in capture probability may significantly affect priming distributions, reflecting sensitivity to immune system–microenvironment interactions. These outcomes highlight a strong relationship between antigen processing efficiency and both the magnitude and pace of immune activation.

Our approach introduces a computational framework that connects neonatal behaviors, specifically superficial scratching, with immunological development, isolating discrete biological events and allowing their probabilities to shape immunological outcomes over time. It merges basic physical phenomena such as cutaneous microtrauma with immune decision thresholds under stochastic variability. By emphasizing time-resolved outputs and varying biological parameters, it enables controlled in silico experimentation, a feature not typically available in purely observational or clinical studies. The model’s primary advantage is its modularity, allowing biological assumptions to be modified and expanded without loss of internal consistency. Because the simulation output is cumulative and time-indexed, the resulting data support detailed kinetic evaluations, enhancing our understanding of early immune response dynamics. In contrast to traditional immunological modeling relying on continuous-time differential equations or population-level compartmental models, our simulation adopts a discrete-time, agent-free stochastic framework that emphasizes minimal assumptions and maximal modularity. While differential equation models can describe the dynamics of known cytokine concentrations or cell populations under fixed initial conditions, they are less effective at capturing the probabilistic and threshold-dependent nature of early immune priming in highly variable individual cases (Liu et al., 2021; Fay et al., 2023; Mallick et al., 2025). Moreover, most existing models do not explicitly consider the skin as an initiator of immune responses in neonates, nor do they incorporate behaviors such as self-scratching as biologically relevant. Most existing models of skin-immune interactions concentrate on chronic inflammatory conditions such as atopic dermatitis or immune responses in adult skin, while largely overlooking the role of acute microtrauma in healthy neonates (Lunjani et al., 2021; Conceição-Silva et al., 2022; Zhang et al. 2022; Huang et al., 2023). Our framework also differs from agent-based models by avoiding spatial or individual-cell tracking, thereby reducing computational complexity while still capturing essential dynamics (Pleyer and Fleck, 2022; Camacho-Gomez et al., 2024; Hardman et al., 2024; Metzcar et al., 2025). It is not a substitute for high-resolution biological simulation, but it serves as a bridge between mechanistic plausibility and theoretical inference.

Several limitations must be acknowledged. Our model simplifies immune activation to a threshold-based additive scoring system, omitting molecular and cellular complexities such as cytokine feedback, cell migration and antigen processing delays. The exclusion of spatial dimensions and individual immune cell behavior reduces biological realism and while it serves computational clarity, it prevents the simulation from representing spatially heterogeneous skin environments. Moreover, we used fixed probabilities for injury, microbial ingress and antigen capture, which may be influenced by external variables such as neonatal microbiota composition, scratching frequency or skin integrity across individuals. The model also assumes independence between time steps unless priming occurs, which omits potential temporal correlations in immune signaling. Only three independent seeds were used to generate statistical comparisons between antigen capture scenarios. While each run involved 1000 trials, a larger number of independently seeded simulations would provide tighter confidence intervals and improved robustness. Lastly, the simulation has not been directly calibrated or validated with empirical data from neonatal cohorts or skin immunology experiments.

Our findings suggest pathways for further research and experimental inquiry. The probabilistic structure could be adapted to integrate additional variables such as neonatal skin hydration, microbiome diversity or presence of skin barrier–enhancing treatments. The simulation can also be extended to incorporate immune memory, regulatory feedback or antigen-specific dynamics, creating a more elaborate view of imprinting processes. One potential research direction includes validating the threshold values for immune priming in laboratory models of neonatal skin inflammation, where defined injuries are paired with microbial exposure and local immune markers are tracked. The model also enables formulation of testable hypotheses, such as whether controlled superficial skin stimulation can increase early-life immune diversity without adverse effects. In clinical contexts, investigations could be designed to assess correlations between early skin barrier disruptions and later immunological profiles, such as vaccine responsiveness or allergy risk. Still, population-level modeling using distributions of individual parameters may allow translation from individual simulations to public health inferences.

In conclusion, we addressed the hypothesis that neonatal skin microtrauma, through mechanisms of antigen exposure and immune stimulation, can contribute to early immune priming in a quantifiable and temporally structured manner. Our main theoretical finding is that increased antigen capture probabilities may produce both faster and more consistent immune activation following cutaneous injury in neonates.

## DECLARATIONS

### Ethics approval and consent to participate

This research does not contain any studies with human participants or animals performed by the Author.

### Consent for publication

The Author transfers all copyright ownership, in the event the work is published. The undersigned author warrants that the article is original, does not infringe on any copyright or other proprietary right of any third part, is not under consideration by another journal and has not been previously published.

### Availability of data and materials

All data and materials generated or analyzed during this study are included in the manuscript. The Author had full access to all the data in the study and took responsibility for the integrity of the data and the accuracy of the data analysis.

## Acknowledgements

none.

## Authors’ contributions

The Author performed: study concept and design, acquisition of data, analysis and interpretation of data, drafting of the manuscript, critical revision of the manuscript for important intellectual content, statistical analysis, obtained funding, administrative, technical and material support, study supervision.

## Declaration of generative AI and AI-assisted technologies in the writing process

During the preparation of this work, the author used ChatGPT 4o to assist with data analysis and manuscript drafting and to improve spelling, grammar and general editing. After using this tool, the author reviewed and edited the content as needed, taking full responsibility for the content of the publication.

